# Scale-dependent predictability of microscopic brain structure from MRI using paired PS-OCT

**DOI:** 10.64898/2026.11.25.769330

**Authors:** Huangjing Ni, Huisheng Zhou, Wentao Jiang, Ming Song

## Abstract

Cortical layering, fascicle geometry and tissue texture of the mouse brain lie in the mesoscopic range. That range sits below the hundred-micrometre sampling of a whole-brain magnetic resonance (MR) volume and above the micrometre structure of individual fibre bundles. Polarization-sensitive optical coherence tomography (PS-OCT) reaches into it, and MRI does not. Super-resolution models can enlarge an MR volume to any grid, but the grid size does not decide what the volume contains. We take PS-OCT at 10 µm isotropic sampling as the target and ask whether a 100 µm MRI volume still carries information about each target band. At each candidate magnification we ask whether the source still determines something about the target band. Two readouts on a shared band index answer it. The bands come from an isotropic spherical-shell decomposition. The energy readout predicts a log-RMS energy envelope, which needs no unobservable phase, and a structural-tensor comparison separates geometric from intensity guidance, with the absolute cosine read against an analytic level of 0.50. On nine paired mouse brains the two readouts do not weaken together. The median absolute cosine falls from 0.864 at 1x to 0.661 at 4x and to 0.518 at 10x, while the *R*^2^ of the energy envelope decays from 0.627 to 0.438 at 4x and to 0.175 at 10x. The two readouts part company at 4x, a target voxel of 25 µm, the last of the six candidates at which both conditions hold. Each candidate interrogates a different physical scale. The curves therefore cannot carry a claim about a special frequency, and we report the boundary as the output of a procedure and not as a constant. Two further effects pull the readouts in opposite ways. One is intrinsic to the envelope kernel and accounts for part of the accelerated decay beyond 4x. The other comes from the smoothing applied to the reference before it is scored.

## 1 Introduction

Magnetic resonance imaging (MRI) delivers whole-brain coverage of a fixed mouse brain, but the anatomical volume used in cross-scale work is sampled at roughly 100 µm isotropic, and structures below that size are not represented. PS-OCT resolves the same brain considerably further: reconstructed whole-brain volumes are represented on a 10 µm isotropic grid [1]. Between the two lies a range in which cortical layering and fascicle geometry become visible, and where a label-free optical contrast is available that MRI does not supply [2, 3]. A method that added PS-OCT-like detail to an already acquired MRI volume would therefore reach structures the MR volume never recorded. Whether it can do so turns on a question about the data, not about the model. Given a source volume sampled at 100 µm, is there a magnification at which the source still carries information about the finer band it is asked to supply? Or does the request pass beyond what the sampling mechanism ever recorded?

That question has been approached from several directions, and each settled a different part of it. The first settled whether the two modalities can be brought into agreement on the same specimen. Serial optical coherence scanning was cross-validated against diffusion tensor imaging in the human medulla [4]. PS-OCT has been mapped onto ultra-high-field diffusion MRI in the macaque brain [5], compared with tractography in the human brainstem [6], and bridged to white-matter geometry across anatomical scales and species [7]. Agreement was reached, usually for a reconstructed fibre angle, at the one scale each study chose to validate. The same work also shows the cost of the reference. Micrometre-resolution whole-brain mapping exists for human hemispheres, non-human primates [8] and the developing human brain [9], and it is confined to a few laboratories. A second direction asked which way the relation runs, and the dataset we build on is an example. Regionally, T2-weighted signal associates with PS-OCT intensity, and more closely with retardance, which the authors read as covariation within a specimen rather than as a registration check or an equivalence of modalities [1]. The relation is therefore real but regional, and it says nothing yet about whether it survives at finer scales. A third direction asked how such a relation can be turned into a volume. Learned image-to-image translation showed that a mapping between two domains can be recovered from paired examples [10], and cycle consistency removed the need for pairs [11]. The medical variants now run from CT to MR and MR to CT synthesis, with diffusion models supplying the generative step in the most recent designs [12, 13, 14]. OCT has been super-resolved within its own modality [15], and surveys order these designs by architectural family [16, 17, 18, 19]. Appearance, however, is what a learned map reproduces, and reproducing appearance is not the same as recovering structure. A fourth direction questioned the magnification itself. Single-image super-resolution has treated it as a free parameter since the move from regression through perceptual and adversarial objectives [20, 21] to diffusion-based and implicit-representation formulations [22, 23], a trajectory the surveys of the field record [24]. Frequency entered that literature as a target rather than as a measuring rod. Frequency terms penalise the spectral distortion that pixel losses under-weight [25], and frequency-guided conditioning couples a frequency branch to a spatial one [26]. That flexibility belongs to the output representation. It does not answer whether the source supports the band being requested, a distinction argued explicitly for super-resolution in microscopy [27]. The four directions share one premise: the target scale stays fixed, because a fixed scale is what makes registration and validation tractable. None of them reports how the agreement behaves as that scale is refined. Once a method is deployed, the premise gives way: a user who asks for a grid finer than the calibration scale is extrapolating away from the only scale at which the relation was checked. No existing result says how quickly the check stops applying. What is missing, then, is the coupling read as a function of the target band: how much of the band’s information survives in the source at each candidate scale, rather than whether the two modalities agree at the scale that was tested.

In this work we measure that coupling rather than infer it from model output. A probe built on a single band index computes two readouts on the same bands, one from tensor orientations and one from band energy, and reads the first against an analytic isotropic level. The two readouts do not weaken together. Orientation holds well beyond the scale at which the energy envelope has already lost most of its explanatory power. The two cross at a single candidate, 4x, a target voxel of 25 µm, which is the last at which both conditions still hold. Past that candidate the source no longer supports one of them, so whatever detail appears at the finer bands is produced and not recovered. Because the curve measures how quickly detail becomes unpredictable rather than locating a frequency at which information stops, the boundary is a property of this source volume and this procedure, not a constant of the modality pair. The Results take up what separates the two readouts and how far each one carries.

## 2 Materials and Methods

The measurement runs as one chain, and the chain separates what belongs to the source from what belongs to the target. The target volume supplies the band structure and the reference values, the source volume supplies every quantity that enters a prediction, and a mapping frozen before any band analysis joins the two, so that no target pixel reaches the source pathway afterwards. A shell decomposition then builds a single band index, and two readouts are computed on it: one from tensor directions and one from band energy. The reference level and the voxel threshold define what a readout is compared against, and the criterion turns the two comparisons into a boundary.

### 2.1 Samples

A band readout compares two volumes of the same brain, so it needs a pair in which the anatomy has been carried across modalities without being selected by anything the readout measures. The rest of this section presupposes such a pair. The specimens are adult male C57BL/6J mice aged 12 to 14 weeks. They were perfusion-fixed with 4% paraformaldehyde after transcardial perfusion under pentobarbital sodium anaesthesia at a dose of 70 mg per kg body weight, and embedded in 10% gelatin for ex vivo imaging. Tissue preparation followed the dataset we build on [1], and the acquisition parameters of each modality are restated in the sections that use them, so that the probe can be reproduced from this paper.

An animal enters the cohort only when a whole-brain PS-OCT reconstruction and a matching ex vivo T2-weighted MRI volume are both available, so the nine pairs are the animals that carried both modalities; no animal was selected or discarded on the basis of any band readout or any other measurement reported here. The mouse is the biological unit of the cohort and of every leave-one-subject-out fold, and the voxel is the unit of the readouts, taken as a patch drawn inside the parenchyma of one subject and bound to that subject so that a held-out brain contributes to no fitted quantity. Subject identifiers are carried across modalities unchanged.

The pairing is fixed before either modality is acquired. Each acquisition is then described in the section that uses it, because the source and the target enter the probe in different roles and are processed on their own terms.

The reconstruction grid and the physical resolution are two different quantities, and for the target we keep them apart here rather than in the discussion. The grid is a representation of the reconstructed volume, not a resolving power. Native sampling before reconstruction is about 5.86 *×* 5.86 *×* 3.52 *µ*m^3^, and no point-spread function was measured for it. Throughout this paper, therefore, 10 µm is a spacing on which the target is represented, and no claim is made about the optical instrument’s ability to separate two structures closer than that. The reconstruction described below builds that grid, and we use its intensity channel as the reference.

From these volumes onward every calculation in this paper is ours. The cross-modal registration, the two filters, the band decomposition, the readouts, the reference level and the criterion are not part of the dataset release. All of them are described below, in the order in which they enter the chain.

### 2.2 T2-MRI and data processing

The readouts need a source grid whose voxels are already isotropic, so that a band defined on frequency modulus means the same thing along every axis. The T2-weighted volume provides that grid once it is registered and resampled.

#### Acquisition

The volume was acquired on a 9.4 T Bruker BioSpec 94/30 AV3HD system with an eight-channel rat-head array coil, using a 3D TurboRARE sequence with RARE factor 4, TE = 20 *ms*, TR = 2000 *ms* and four averages over a 120 *×* 110 *×* 170 matrix and a 12 *×* 11 *×* 17 *mm*^3^ field of view, giving 100 µm isotropic sampling. No gadolinium-based staining was used.

#### Processing

The alignment chain is rigid, affine and symmetric diffeomorphic [28], implemented in ANTs and driven by Mattes mutual information and neighbourhood cross-correlation on the two macro-scale outlines. The deformation field was then frozen. Intensities were truncated at the 1st and 99.9th percentiles per subject and mapped linearly to [*−*1, 1], with background set to *−*1. Each candidate is resampled onto its own grid with a third-order three-dimensional Butterworth low-pass, whose 3 dB cut-off sits at 5.0 cycles/mm and whose attenuation is *−*24.2 dB at 10 cycles/mm. That filter suppresses residual energy at and above the 4x target band before any analysis, and its cut-off is held at this value for every candidate. Every band readout is computed on 64^3^ patches selected inside the brain parenchyma.

This section fixes the standing of the source in everything that follows. The T2-weighted volume is the only input any readout receives, and the frozen deformation means that every source-side quantity, including the nine-dimensional context vector of the prediction protocol, is a function of this volume alone. In that sense the source is fixed before the first band is formed, and the same fact is why the target volume can define bands and score readouts without leaking into the prediction.

### 2.3 PS-OCT imaging and data processing

The target has to supply two quantities that differ in kind, a direction that can be compared with the source’s direction and an energy that can be compared with the source’s energy, and both have to come from the same acquisition. PS-OCT supplies them because a spectral-domain measurement returns the amplitude and phase of light backscattered from each depth, and birefringence along the optic axis turns the phase difference between two polarization channels into a direction [2, 3]. A modality that carried only intensity would supply the energy and not the direction, and one that carried only retardance would supply the direction and not a phase-invariant energy. That is why the reference here cannot be replaced by a histological stain or a bright-field section.

#### Acquisition

Volumes were acquired with a Thorlabs Telesto polarization-sensitive spectral-domain system at a nominal centre wavelength of about 1300 nm and an A-scan rate of 28 kHz, with tiles of 512 *×* 512 *×* 256 voxels at 5.86 *×* 5.86 *×* 3.52 *µ*m^3^ and a 3 *×* 3 mm^2^ lateral field, staged on a 2.4 mm grid with about 20% overlap in a typical 4 *×* 5 tile array per level.

#### Processing

Tiles were registered by phase correlation on the intensity channel and fused, and levels were aligned and combined to whole-brain intensity and retardance volumes on a 10 µm isotropic grid, by the procedure of the dataset release [1]. As noted above, the grid is a spacing and not a resolving power. The energy readout is scored against the intensity channel of the reference. The retardance channel is not used, because the readout it would enter is the geometric one, and that readout needs a direction rather than a contrast. One further filter belongs to this side of the measurement, at the point where the verification model scores the reference. The 10 µm reference is smoothed with a separable Gaussian whose standard deviation is one half of the target voxel, that is 1.25 voxels of the reference grid or 12.5 µm, on a 5 *×* 5 *×* 5 support, before trilinear resampling onto the 25 µm grid. The two filters acting in this study, the Butterworth resampling of the source and this smoothing of the reference, are not interchangeable. A reader who takes them for one would attribute to the coupling what one of them removed. What this width does to the upper part of the 4x band is given with the results, and the discussion returns to what it does to the reported *R*^2^. Two properties of the target follow from this construction, and neither is a defect of it. The reference carries the sectioning marks and the level-to-level intensity steps that the reconstruction could not remove entirely, and those features sit in the bands the readouts score. The verification model smooths rather than reproduces them, which is what a model working inside a boundary should do. In addition, the finest candidates on the enumeration are the ones closest to this grid’s own Nyquist limit, and that is why the band index is restricted away from that edge.

### 2.4 Isotropic shell decomposition

The split between the two registered volumes leaves no coordinate on which a band can be named, and both volumes need a shared coordinate on which “the same band” means the same thing at every candidate. A cubical band in the three frequency axes cannot supply it, and the reason is geometric rather than practical. A cube in (*f_x_, f_y_, f_z_*) does not correspond to a physical length scale, so an oblique frequency is assigned to a band it does not belong to. Because the volumes are isotropic, the modulus of the spatial frequency is the correct coordinate, and concentric spherical shells follow from that choice rather than being imposed on it.

Each reference patch is tapered with a separable three-dimensional Hann window on all three axes before the transform. The taper removes the discontinuity at the patch boundary, which would otherwise appear as a cross-shaped artefact in the spectrum. Writing **f** = (*f_x_, f_y_, f_z_*) in cycles per millimetre, the shell mask for band *k* is

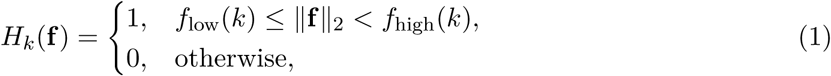

with the integration range restricted to [0.2 *f*_Nyq_(*k*), 0.8 *f*_Nyq_(*k*)] for each band, so that neither the Gibbs ringing at the discrete Nyquist limit nor the slowly varying bias near zero frequency is attributed to a shell. Inverse transformation of *H_k_I*^^^ gives the band-limited field *I_k_*(**x**). This quantity is the input to both readouts that follow.

The size of the working aperture follows from the transform at its own edges, not from a preference about band widths. Below 0.2 *f*_Nyq_ the slowly varying bias of the taper dominates what the shell contains, and above 0.8 *f*_Nyq_ the Gibbs ringing of the finite support does. The window is therefore placed between those two fractions of Nyquist for every band. Inside the window, the candidate fixes the band edges, and Table 1 lists them.

**Table 1:** The six candidate magnifications and the spatial-frequency band each one targets. Period is 1000*/f*, the structural size the band interrogates. *f*_norm_ is the band expressed as a fraction of 20 cycles/mm, the Nyquist frequency of the 25 µm grid; *f*_own_ is the same band expressed as a fraction of the Nyquist frequency of the grid on which that candidate is evaluated.

| Mag. | Target (µm) | Band (cycles/mm) | Period (µm) | *f*<sub>norm</sub> (25 µm) | *f*<sub>own</sub> |
|---|---|---|---|---|---|
| 1x | 100 | 2 to 5 | 200 to 500 | 0.10 to 0.25 | 0.40 to 1.00 |
| 2x | 50 | 5 to 10 | 100 to 200 | 0.25 to 0.50 | 0.50 to 1.00 |
| 4x | 25 | 10 to 20 | 50 to 100 | 0.50 to 1.00 | 0.50 to 1.00 |
| 6.25x | 16 | 15 to 31.25 | 32 to 67 | 0.75 to 1.56 | 0.48 to 1.00 |
| 8x | 12.5 | 20 to 40 | 25 to 50 | 1.00 to 2.00 | 0.50 to 1.00 |
| 10x | 10 | 25 to 50 | 20 to 40 | 1.25 to 2.50 | 0.50 to 1.00 |

Table 1 gives the six candidate magnifications, the physical band each one targets, the period that band corresponds to, and the two normalisations of the band tested against the coordinate-system question. Table 3 reports the measured values obtained at each of them. The six magnifications are the ones that the reconstruction and resampling chain supports exactly. The criterion itself is defined on a continuous frequency axis, so the spacing of the candidates is a property of the study rather than of the criterion.

### 2.5 Energy envelope as a phase-invariant target

The first readout has to say how much of the target band’s energy the source still explains. The band-limited field *I_k_* cannot serve as the quantity to be predicted, because its absolute phase at fine scales is not identifiable across individuals. What is stable is how much anisotropic scattering occurs in a neighbourhood, and the envelope is built for that quantity rather than for the field. Two alternatives fail here. A pointwise regression on *I_k_* has no stable minimiser other than the conditional mean, since a pointwise loss asks for a value the phase does not provide. A texture statistic such as a local gradient magnitude responds to the arrangement rather than to the content of the band, and it rises when a band is filled with interpolated structure.

We therefore predict the local energy of each band rather than its phase,

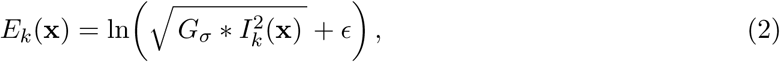

where *G_σ_* is a separable Gaussian of standard deviation one voxel on a 5 *×* 5 *×* 5 support, evaluated on the 10 µm reference grid. The logarithm compresses a dynamic range that spans several orders of magnitude and turns a multiplicative relation into an additive one that linear regression can express. This quantity is then the target of the prediction protocol described next.

The standard deviation of the kernel is set by what the envelope has to remain sensitive to, not chosen for convenience. Squaring the band-limited field and then smoothing it applies the kernel to the squared signal, so the envelope responds to a band through the square of the Gaussian transfer function. A narrower kernel would place the half-amplitude point proportionally higher and would therefore follow more of the finest bands, at the price of a noisier estimate of local scattering. A wider kernel would be steadier and would leave less of every band inside the passband. The width is therefore a trade between the noise of the estimate and the part of a band the estimate is still sensitive to, and the results contain the passband that follows from this width.

Two things follow, and they are worth separating. The envelope is a well-posed regression target because it discards exactly the quantity that is not identifiable. It is not a description of texture: a method could predict *E_k_* accurately and still fail to synthesise plausible microstructure, because the envelope fixes the local intensity of scattering but not its arrangement. The verification model shows that the arrangement is at least not collapsed, and no more than that.

### 2.6 Joint structural tensor and the isotropic null

The second readout has to answer a question the energy readout cannot: whether the direction of structure at the target scale is still informed by the direction at the source scale. The envelope is blind to it because the envelope discards phase. A cubical region in which a single dominant direction is assumed would not do either, since the tissue it is being applied to contains boundaries and isotropic patches. A structure tensor tolerates both, because its principal eigenvector is defined even where the eigenvalue contrast is small, and its eigenvalue spread says how far that direction can be trusted. That estimator is the one standard for serial optical coherence data [29], and its accuracy in brain-tissue microscopy depends on the window scale and on the noise level, both of which are held fixed across the two modalities here [30]. We compute the same tensor on both volumes with the same differential Gaussian kernel and no modality-specific diffusion parameters, and take the principal eigenvector at each voxel as the local tangent direction. The macro-scale tangent on the MRI side is compared with the micro-scale direction recovered from PS-OCT through

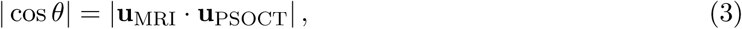

where the absolute value removes the sign ambiguity that a tangent direction carries.

The comparison needs a baseline, and we state it here because a readout without one cannot be read at all. A median absolute cosine carries no meaning on its own until one knows what the same statistic would be if the source carried no directional information about the target. Consider two independent directions drawn uniformly on a hemisphere, paired without regard to which belongs to which specimen. Writing *θ* for the angle between them, an isotropic axis has a component along a fixed direction that is itself uniform, so cos *θ* is uniform on [*−*1, 1] and its absolute value is uniform on [0, 1]:

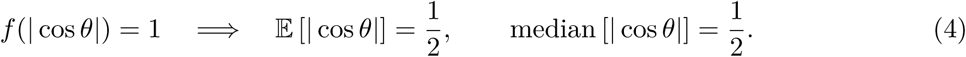

The mean and the median therefore coincide, and either can be used as the reference level. We work with the mean, 0.50, because it is the value a specimen would average to if its macro-scale orientation carried no information about its micro-scale orientation, and because the midpoint of a uniform distribution is the least parameterised of the available summaries.

That level is the weakest admissible baseline, and its weakness is part of the argument. Because the tissue is not isotropic, a steeper null follows from breaking the pairing between specimens, and such a permutation inherits the anisotropy, the tensor window and the patch selection of the actual data instead of assuming them away. The permutation keeps every estimated tensor and re-pairs the patches across subjects, so that a patch from one brain is compared with a patch from another at the same registered anatomical location and in the same band; the statistic is the same median absolute cosine. Because the tensors themselves are unchanged, the permutation isolates the contribution of subject-specific correspondence. Because the location is matched through the frozen registration, it holds the macro-scale geometry fixed. Exceeding 0.50 is therefore necessary rather than sufficient, and the excess of a measured median over that level is the quantity a stronger null would have to be tested against. Figure 1 summarises the geometry behind the reference level: under isotropic directions the absolute cosine is uniform, so the stated 0.50 is where that distribution is centred.

**Figure 1:**
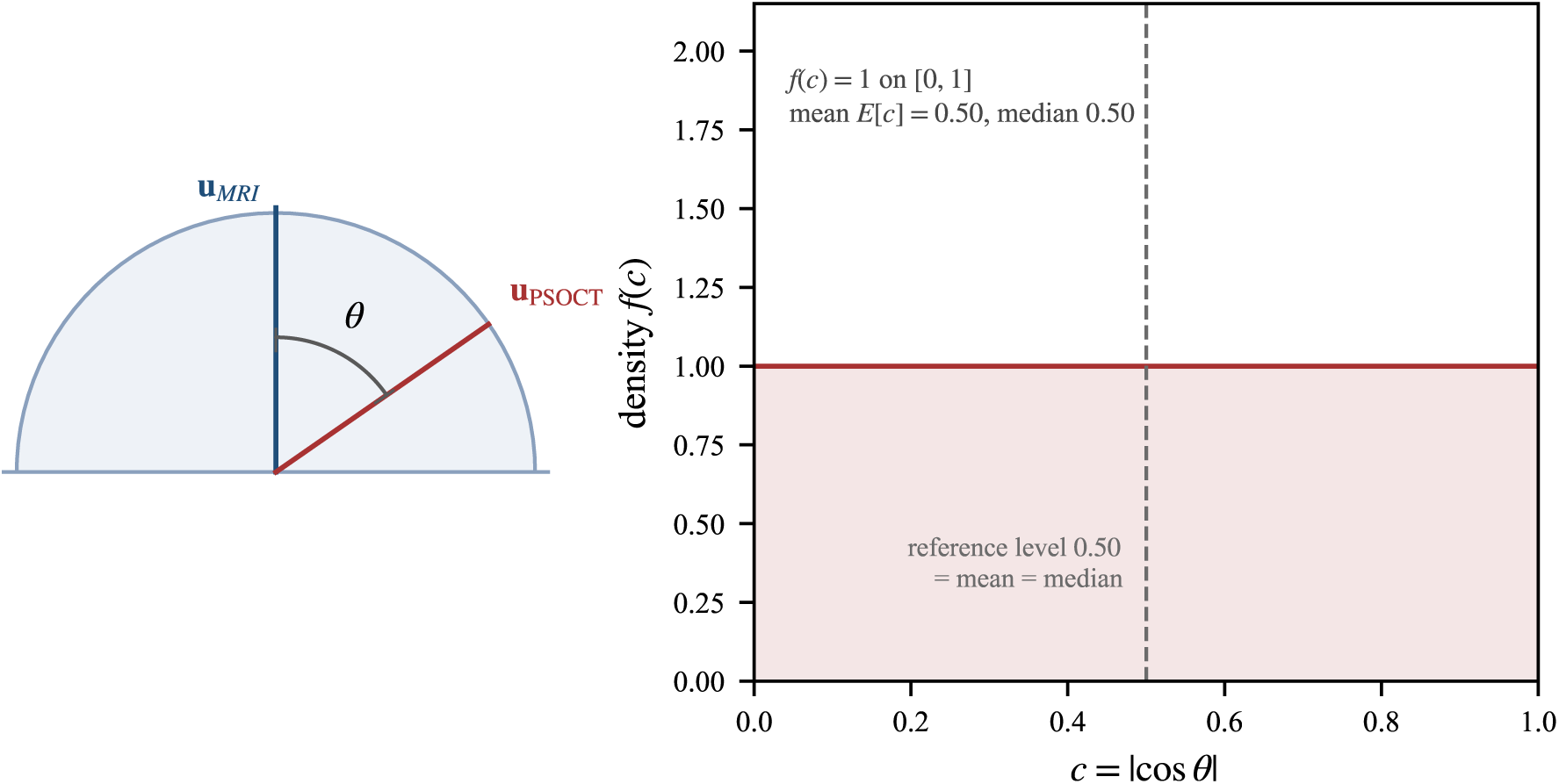
Geometry of the isotropic reference. Left: two independent directions drawn uniformly on a hemisphere, with *θ* the angle between them. Right: the induced distribution of the absolute cosine, which is uniform on [0, 1], so that its mean and its median are both 0.50. The reference level is that midpoint.

One threshold class remains in this section, and its value and its fixity both need a reason. Because the tensor estimate is dominated by boundaries in some regions and is ill-conditioned in isotropic ones, the median is taken only over voxels whose fractional anisotropy exceeds 0.3, computed from the three eigenvalues of the same tensor. We fixed the threshold once and applied it identically in all six bands and in both modalities. The threshold is permissive, so that the filter removes voxels in which no principal direction is defined rather than narrowing the median to the most strongly oriented tissue; what the cut admits, in eigenvalue terms, is given with the results. The results also report, band by band, how much tissue survives the cut, and that population enters the reading of the curve. This section reports the level and the cut that every median in this paper depends on, and it stops there; how a measured median is compared with that level belongs to the rule stated below.

### 2.7 Leave-subject-out prediction ceiling

The energy readout has a target, *E_k_*, and this section fixes how much of it a source-side predictor can be credited with. The split between source and target does not settle that by itself, because a fit that has seen voxels from the brain it is scored on reports its own interpolation as predictability.

The source-side features are the MR intensities and their local statistics at several spatial scales, forming a nine-dimensional context vector per voxel, and we compute every feature on the MRI side of the frozen mapping. For each band, a ridge regression on these features predicts the log-RMS envelope. The protocol is leave-one-subject-out: each fold holds out one whole brain, fits on the remaining eight, and evaluates *R*^2^ = 1 *−* (*y − ŷ*)^2^*/* (*y − ȳ*)^2^ on the held-out volume only, so that spatially correlated voxels from the same brain cannot contribute to their own prediction. Holding out a whole brain rather than a voxel block is what makes the score a statement about a brain the predictor has never seen, and that is the sense in which the number can be compared across candidates at all. We set the ridge penalty inside each fold rather than for the whole study, so that the strength of the penalty cannot be tuned on the volume that is being scored.

The quantity this protocol returns is therefore a ceiling on what a linear predictor reading the source can explain about the target band’s energy. It is not an estimate of a causal relation, and it is not a measure of how much structure a generator will recover. The predictor is also deliberately simple: a more expressive one would raise the curve, so the shape of the curve belongs to the coupling rather than to the capacity of the predictor. The criterion below compares this ceiling against its floor, and the results describe how it varies across the six candidates.

### 2.8 Boundary criterion and its margins

The sections before this one deliver two scores and one reference level, and none of them is a boundary. A boundary has to be defined rather than read off a curve, and the definition used here is the following.

The first step fixes the reference modality and its grid, then resamples the source onto each candidate grid with an anti-aliasing filter whose stop band begins at the finest target band under consideration. The source then cannot carry residual energy at a band it is later credited with informing. Each candidate is resampled and decomposed on its own grid, and a normalisation computed for one grid is not carried over to another; carrying it over would turn the rejection at the fine end into a statement about the grids rather than about the modalities. The candidate magnifications are then enumerated on the continuous frequency axis, each with its band, the physical period that band corresponds to, and its normalised frequency, as in Table 1. Because the axis is continuous, the enumeration cannot be arranged around a preferred answer, and the spacing of the six candidates is a property of the study rather than of the rule.

Once the bands exist, the two readouts are computed on them, the median absolute cosine and the leave-subject-out coefficient of determination. A readout computed with its own refit on the brain it is scored on would report its interpolation as coupling, which is why the leave-one-subject-out protocol is part of the rule rather than a refinement of it. The scores are then read against a null, taken either from the geometry of the tensor comparison or from a permutation of subject pairing, with the analytic isotropic value as the weakest admissible baseline. A baseline steeper than the data justify would turn a real excess into a null result, so the strength of the baseline is itself part of what has to be stated.

We locate the boundary at the largest candidate at which the geometric readout exceeds the null by a stated margin while the energy readout exceeds its floor by a stated margin. The first candidate at which either condition fails is reported as the point past which the boundary lies. Naming the boundary after the first candidate that satisfies both conditions would move the answer to the earliest candidate that is admitted at all, which answers how early the source begins to weaken and not how far it can be pushed. Two further steps follow from the same concern. The sensitivity of that location to both margins is reported as the range over which the answer does not move, because a boundary quoted as a number while its margins were never varied is not a located boundary. What the normalisation does to the answer is reported as well, together with whether accepted and rejected candidates differ in their normalised position or only in their measured readouts, since an answer whose position changes with the coordinate system belongs to the grids and not to the modalities. Both steps are measurements and not parts of the rule, and the results report them.

Three properties of this rule decide where it can be carried. Its inputs are paired images of the two modalities, a registration that can be frozen, and a null that can be computed from the geometry or from a permutation. Any macro-to-micro pair that supplies those three can be treated the same way, and the candidates we have in mind are polarized light imaging, bright-field histology and synchrotron phase contrast, each of which serves as a structural reference for an anatomical volume. The only requirement on such a reference is that it carry both an orientation-bearing quantity and a phase-invariant energy. What the rule returns is a frequency, not a class label, so a method operating inside the boundary is not certified as faithful; it is certified as operating in a band where the source still constrains something. The last step of the account above is what keeps the answer from depending on the coordinate system. Two normalisations can always be found that disagree, and a boundary whose position moves between them is a boundary about the grids and not about the modalities.

### 2.9 Evaluation metrics and reporting conventions

The band readouts need no external metric, because a geometric reference or a held-out fit defines each of them. The feasibility check scores a synthesised volume, and a score of that kind is defined by a published formulation. We therefore name the three formulations used here and add nothing to them. Structural similarity follows its standard definition and its multiscale variant as used in image-quality studies [31, 32]. Learned perceptual distance follows the LPIPS formulation [33]; the deep feature backbone that formulation uses is a pretrained convolutional image classifier [34], and its identity is part of the metric, because the value depends on which network supplies the features. We compute the perceptual score on axial, coronal and sagittal slices taken at five-voxel spacing and aggregate it by tissue-voxel weight. Peak signal-to-noise ratio needs no reference. All three are computed on parenchymal voxels of the union brain mask, over the nine leave-one-subject-out folds, so that every score reported for this check is a mean with its standard deviation across held-out brains. Each score is reported with the dispersion across folds and is read against the band analysis rather than against a baseline.

## 3 Results

### 3.1 Orientation survives the mid-range bands, then fails

We compute the geometric readout on voxels whose anisotropy clears the 0.3 cut fixed for the tensor comparison, and take it as the median absolute cosine over the same patch population used for the energy readout. In eigenvalue terms that cut is permissive and not selective: for a tensor whose two minor eigenvalues are equal, an anisotropy of 0.3 corresponds to a leading eigenvalue about 1.65 times the others, so the cut excludes voxels in which no principal direction is defined and leaves the weakly oriented tissue in the median. On nine paired brains the median has a definite shape, and the shape is not monotone. In the lowest band, at a period of 200 to 500 µm, it is 0.864, an angle of 30.2*^◦^*, far from the isotropic level and consistent with a macro-scale boundary guiding the direction of the micro-scale fibre bundles it encloses. At 2x it falls to 0.640, an angle of 50.2*^◦^*. At 4x it is 0.661, an angle of 48.6*^◦^*, which is slightly above the value one candidate earlier.

That small rise is smaller than any between-subject dispersion we could plausibly attach to a statistic computed on a selected subset of voxels. The gap is 0.021 in the median and no interval estimate accompanies it, so we do not read the rise from 2x to 4x as evidence that macro-scale guidance strengthens between those two bands. That would be a claim about coupling, and the data do not support it. What the curve does support is the contrast between its ends, where the gaps are large: the readout moves from 0.864, where a macro-scale direction clearly informs a micro-scale one, to 0.518, where any such information is at the level of noise.

The dip at 2x has an explanation that the survival figures support, and the explanation belongs to the voxel population rather than to the tissue. The share of parenchymal voxels passing the anisotropy cut falls from 98.5 per cent at 1x to 96.0 at 2x, to 89.1 at 4x, and to 78.3 at 10x (Table 2). A band that admits a larger share of weakly anisotropic voxels shows a lower median, because weakly oriented voxels contribute near-null directions to the pooled sample. The step from 2x to 4x is the case in point: the admitted share drops by 6.9 percentage points, roughly 4.7 times the between-subject standard deviation of either band, and the median rises by 0.021, which is the size of rise the shrinkage of the admitted population predicts. The dip is therefore a consequence of the changing voxel population and not an unexplained feature of the curve.

**Table 2:** Share of parenchymal voxels passing the anisotropy threshold in each band. Pass is that share as a percentage; SD is its between-subject standard deviation in percentage points; Range is the smallest to largest per-subject value, in per cent.

| Mag. | Target (µm) | Band (cycles/mm) | Pass (%) | SD | Range (%) |
|---|---|---|---|---|---|
| 1x | 100 | 2 to 5 | 98.5 | 1.56 | 94.9 to 100.0 |
| 2x | 50 | 5 to 10 | 96.0 | 1.47 | 93.2 to 98.7 |
| 4x | 25 | 10 to 20 | 89.1 | 1.39 | 85.7 to 90.2 |
| 6.25x | 16 | 15 to 31.25 | 82.0 | 1.93 | 79.5 to 85.2 |
| 8x | 12.5 | 20 to 40 | 79.3 | 2.23 | 75.8 to 82.5 |
| 10x | 10 | 25 to 50 | 78.3 | 1.96 | 74.1 to 80.6 |

The same account does not carry past 4x. Beyond that candidate the readout declines to 0.553 at 6.25x, to 0.530 at 8x and to 0.518 at 10x, where the median angle is 58.8*^◦^* and the median exceeds the isotropic mean of 0.50 by 0.018. The admitted share keeps shrinking over the same four candidates, and a shrinking admitted population would push the median up. It does not, so whatever drives the decline there acts on the tissue rather than on the selection, and the weakening exceeds what the population shift can offset. The excess over the isotropic level at that candidate is 0.018, and we do not treat it as guidance.

Of the two quantities a band can carry, then, direction is the one the source keeps longer. At 10x the median still exceeds the isotropic mean of 0.50 by 0.018, so directional agreement is not extinguished at the finest candidate, although it has fallen close to the level of noise. How far the energy carried by the same band survives is a separate question, and the next subsection measures it.

### 3.2 Energy stays predictable only inside the envelope passband

The energy readout is the leave-subject-out coefficient of determination of the log-RMS envelope, so the score depends on a residual that the held-out brain never touched, and the curve it produces has no reversal (Figure 2). It falls from 0.627 at 1x to 0.553 at 2x, 0.438 at 4x, 0.327 at 6.25x, 0.257 at 8x and 0.175 at 10x. Read as a sequence, the values describe a monotone loss of predictability rather than a cliff. That shape already decides how the boundary may be described: with no single drop to point at, a boundary stated as the place where the curve falls would be arbitrary.

**Figure 2:**
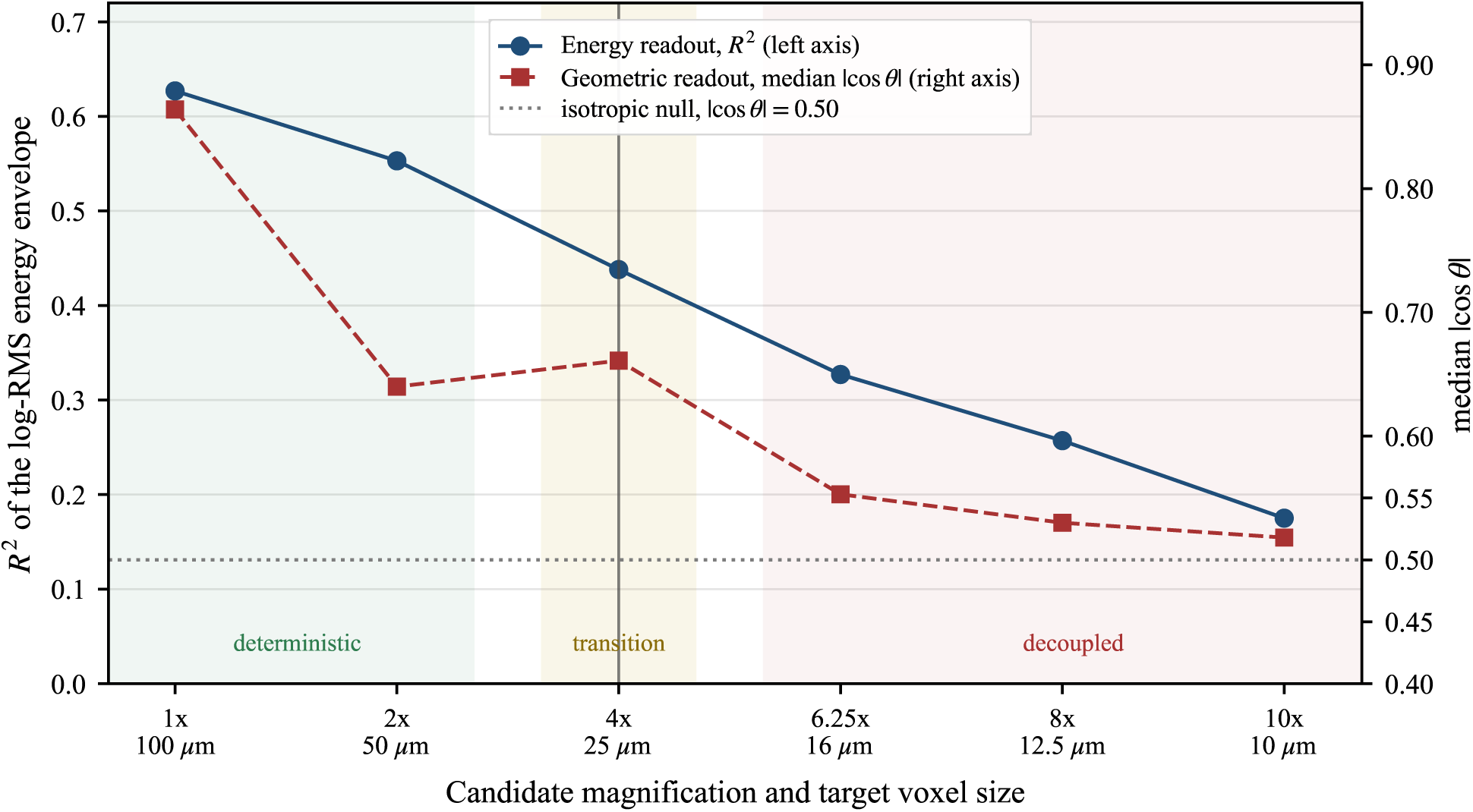
Both readouts as a function of the candidate magnification, labelled with the target voxel size of each candidate. The energy readout is a coefficient of determination against the left axis; the geometric readout is a median absolute cosine against the right axis, with the analytic isotropic level of 0.50 drawn as a dotted line. A vertical line marks 4x. The background bands are a visual guide only, and no part of the argument depends on them: it rests on the six measured values listed in Table 1.

Some of the acceleration beyond 4x belongs to the envelope kernel, not to the tissue. That kernel is a Gaussian applied to the squared band-limited field, so the square of its transfer function gives its effect on a band. For one voxel on a 10 µm grid the envelope response falls to half amplitude at 13.25 cycles/mm, to one tenth at 24.15 cycles/mm and to one hundredth at 34.15 cycles/mm. The 4x band runs from 10 to 20 cycles per millimetre, so the fraction of it whose envelope response is at least one half is one third. For 6.25x, whose band begins at 15 cycles per millimetre, the fraction is zero, and it stays zero for 8x and 10x. At the three finest candidates, therefore, the energy readout is computed on a target field from which most or all of the requested band has been removed, and a decline there is expected even if the underlying coupling were unchanged.

This reading is falsifiable without new image data or new animals. Recomputing the envelope with a narrower kernel, one half voxel rather than one, would move the half-amplitude point from 13.25 to 26.50 cycles/mm and bring the whole of the 4x band and 71% of the 6.25x band inside the passband. The account predicts that *R*^2^ at 4x would then rise by more than it rises at 6.25x, because the kernel is the stronger constraint on the coarser of the two bands. A rise of equal size at both candidates would instead indicate that the kernel is not the limiting factor, and that the decay reported here is a property of the coupling. The same reasoning also sets an upper limit on how far the envelope readout can be trusted: a kernel wide enough to be a stable estimator is a kernel narrow enough to score only the lower part of the finest bands.

Direction and energy therefore do not measure one quantity twice. Orientation is estimated from the geometry of the tensor and is insensitive to overall intensity, whereas the energy envelope is a smoothed scalar field whose passband has to be reported alongside its score. A band can retain directional agreement while its energy has become only partially predictable, and the reverse is possible as well. The next subsection places the two curves on one pair of axes and asks where they stop behaving in the same way.

### 3.3 The two readouts part company at 4x

We now apply the rule stated above to the six measured candidates, and what it returns is a location rather than a ranking. Figure 2 shows the two curves on one pair of axes, and the feature that matters is where they stop agreeing. At 1x and 2x the geometric readout is far above the isotropic mean and the energy readout is near its highest value. At 6.25x and beyond the geometric excess over 0.50 has fallen to 0.053 and below, while the energy readout has lost more than half of its explanatory power. Between these regimes lies 4x. There the geometric readout is 0.661, an excess of 0.161 over the isotropic mean, so macro-scale orientation still carries information about micro-scale orientation. The energy readout is 0.438, well below its 0.627 starting value but not empty, since over two fifths of the envelope’s spatial variance is still explained by a nine-dimensional linear function of the source.

That combination is what we mean by a boundary. Where orientation is still informed, the position of structure can be constrained; where the energy envelope is only partly predictable, the intensity of that structure can be approximated statistically but not determined. In a band where both hold, a synthesis method has something to work with. In a band where neither holds, the output is an educated invention rather than a recovered structure. Table 3 collects the measured values behind the two comparisons, one row per candidate.

**Table 3:** The two readouts at each candidate magnification on the nine paired brains. *R*^2^ is the leave-subject-out coefficient of determination of the energy envelope; *|* cos *θ|*is the median absolute cosine between macro-scale and micro-scale principal directions, taken over voxels that pass the anisotropy cut; Angle is the same median expressed in degrees.

| Mag. | Target (µm) | Band (cycles/mm) | *R*<sup>2</sup> | \|cos *θ*\| | Angle (°) |
|---|---|---|---|---|---|
| 1x | 100 | 2 to 5 | 0.627 | 0.864 | 30.2 |
| 2x | 50 | 5 to 10 | 0.553 | 0.640 | 50.2 |
| 4x | 25 | 10 to 20 | 0.438 | 0.661 | 48.6 |
| 6.25x | 16 | 15 to 31.25 | 0.327 | 0.553 | 56.4 |
| 8x | 12.5 | 20 to 40 | 0.257 | 0.530 | 58.0 |
| 10x | 10 | 25 to 50 | 0.175 | 0.518 | 58.8 |

The claim is about which candidate is last, not about which is first. Among the six measured candidates, 4x is the last at which both conditions hold: at 6.25x the geometric excess is 0.053, a margin we cannot test, and the energy readout has fallen by more than half. The six points are point estimates without intervals, so adjacent candidates are not resolved from each other. We therefore state the result at the level that survives that objection: the two readouts part company at 4x, and every candidate beyond it fails at least one condition.

The location just stated is conditional on the pipeline that produced the scores, and the sub-sections that follow test that condition. Recomputing the envelope with a narrower kernel is the test described above, and until it is run the energy curve carries the response of this pipeline along-side the coupling it measures. The transfer function of the reference-side filter sets part of that response. For the Gaussian width used here it falls to one half at 10.60 cycles/mm and to one tenth at 19.32 cycles/mm, so the upper part of the 4x band reaches the evaluation as a partially smoothed target, and the score of 0.438 describes the target as it is represented and not the unsmoothed optical signal. The next subsection takes up what that condition leaves open about the location itself, and the more practical question of what each readout is good for closes the section.

### 3.4 The exclusion holds under either normalisation

Table 1 names bands that can be placed on a common axis in two ways, and this subsection asks whether the answer to the last question depends on which way is chosen. It does not, and the reason is worth setting out, because the two normalisations fail in opposite ways.

The first normalisation divides each band by 20 cycles/mm, the Nyquist frequency of the 25 µm grid, which is the grid at which the application of interest operates. On that scale the six bands run from 0.10 to 2.50 in normalised frequency, and the three finest candidates lie partly or wholly above unity. Read alone, that column would suggest that the finer candidates ask for a frequency the 25 µm grid cannot hold. The suggestion is idle. No candidate is evaluated on that grid: each is resampled onto its own target grid, and the upper edge of each band coincides with that grid’s Nyquist frequency by construction.

The second normalisation divides each band by the Nyquist frequency of its own grid. It matches how the data were produced, and it fails for a different reason: it returns 0.48 to 1.00 on every row. Every candidate occupies the upper half of its own representable range, so the column locates a band on its grid and says nothing that differs between candidates.

Neither column, then, can adjudicate between candidates. What the pair can do is test whether the exclusion of the two finest candidates is an artefact of the choice, and the test comes out negative. Under the first normalisation, rejecting 8x and 10x means rejecting bands whose normalised upper edge is 2.00 and 2.50. Under the second, it means rejecting bands that occupy 0.50 to 1.00 of their own grids, which is the same interval every accepted candidate occupies. No change of normalisation moves either candidate into a range that the accepted candidates do not already share. The exclusion therefore survives both readings, and it survives for the reason the readouts give rather than for the reason the grids give.

What does not survive is any statement about a special frequency. The bands do not interrogate the same structure: the period column of Table 1 runs from 200 to 500 µm at 1x and from 20 to 40 µm at 10x. A fall in *R*^2^ across the six rows therefore mixes two things, namely that the source knows less and less about the target, and that the question being asked is about finer and finer structure. The curves support the statement that detail becomes progressively less predictable, and they do not support a location in frequency at which information stops.

### 3.5 The boundary stays at 4x across most of the margin range

The answer reported above was produced by comparing two readouts with two stated margins. With six measured points, the question of how much of it is the margins can be answered by enumeration rather than by assumption.

The first part of that answer is negative and it is the more important of the two. The geometric margin enters the location of the boundary only through its position relative to 0.161, the excess at 4x. Any margin from 0.018 to 0.161 leaves the answer at 4x: no candidate beyond 4x clears 0.053, and only a margin below 0.018 would begin to admit 10x. The energy threshold behaves in a similar way over a narrower range. Any value from 0.327 to 0.438 admits 4x and excludes 6.25x, so the answer stays at 4x throughout; a threshold below 0.327 admits 6.25x, and one above 0.438 admits no candidate. The two margins are consequently not interchangeable knobs: the energy threshold decides between 4x and 2x, and the geometric margin decides between 4x and the finer candidates. The second part is a warning about naming. If the boundary is read as the first candidate that satisfies both conditions, then any margin pair that admits 2x makes the answer 2x, because 2x is met earlier. The number reported here is therefore the largest admissible candidate, and it coincides with the first turning point only in the narrow sense that 4x is where the two readouts part company. Read as the first admissible candidate, the same data would place the boundary at 2x for every margin pair that admits it. We report the largest admissible candidate because the question asked of the source is how far it can be pushed, not how early it begins to weaken.

Two degenerate corners are worth stating, and we correct both from an earlier reading of these points. With the energy threshold at 0.439, no candidate satisfies both conditions at any geometric margin, because no measured band reaches that level of predictability, and the procedure correctly returns nothing rather than a boundary. The same happens as soon as the energy threshold is set above 0.438 while the geometric margin is left at 0.15 or 0.16. This is not a failure of the probe; it is the consequence of asking for more energy than any measured band carries.

One further property of the enumeration deserves emphasis, because it cuts against the way this result is likely to be quoted. The boundary stays at 4x over the whole rectangle in which the energy threshold lies between 0.327 and 0.438 and the geometric margin lies between 0.018 and 0.161. It leaves that rectangle in only two ways. Lowering the energy threshold below 0.327 admits 6.25x whenever the geometric margin is at most 0.053. Lowering it below 0.257 admits 8x, but only once the margin is at most 0.030. The admissible range of the geometric margin is nine times wider than that of the energy threshold, so the stability of the answer rests mainly on the energy channel. The geometry makes 4x interpretable; the energy makes it a boundary.

Throughout this enumeration the two margins are the only quantities varied. The anisotropy threshold that selects voxels for the tensor comparison is a third parameter. It is fixed at a fractional anisotropy of 0.3, it changes every median, and we hold it fixed here because sweeping it requires re-running the tensor analysis rather than re-reading the table.

### 3.6 The two readouts diagnose the two failure modes

Once the boundary is fixed, each readout names a different way a generator can fail. Hallucination is what a generator does when geometric guidance has failed. Without a directional constraint, the training prior chooses the principal direction of a synthesised structure, and the result can be a fibre that is locally sharp and globally wrong. The mean solution is what happens when the energy readout has decayed while the loss still asks for voxel accuracy. In the band where the envelope is only partly predictable, the minimiser of a pointwise error lies close to the conditional mean, and the texture that survives is whatever the adversarial term happens to sustain.

The two readouts thus give the two failure modes a quantitative address, and that is the practical reason for measuring both rather than one. A method that improves a perceptual score while leaving the orientation readout at its 10x value has improved the appearance of an invention. Whether that address corresponds to a band a generator can actually use is the last question this section asks.

### 3.7 A generator reaches the target at 4x

The subsections above measure what the source and the target share, and this one asks a different question of the same band: whether a generator trained on the source can reach the target there. The answer is a verification rather than a further measurement, and it runs on the same subject-level hold-out, with a conditional generative model trained at the single magnification of 4x and evaluated on brains held out of its fitting. What the check establishes is that the band identified above is not empty. A companion study reports the design, and what we add here are the scores and Figure 3.

**Figure 3:**
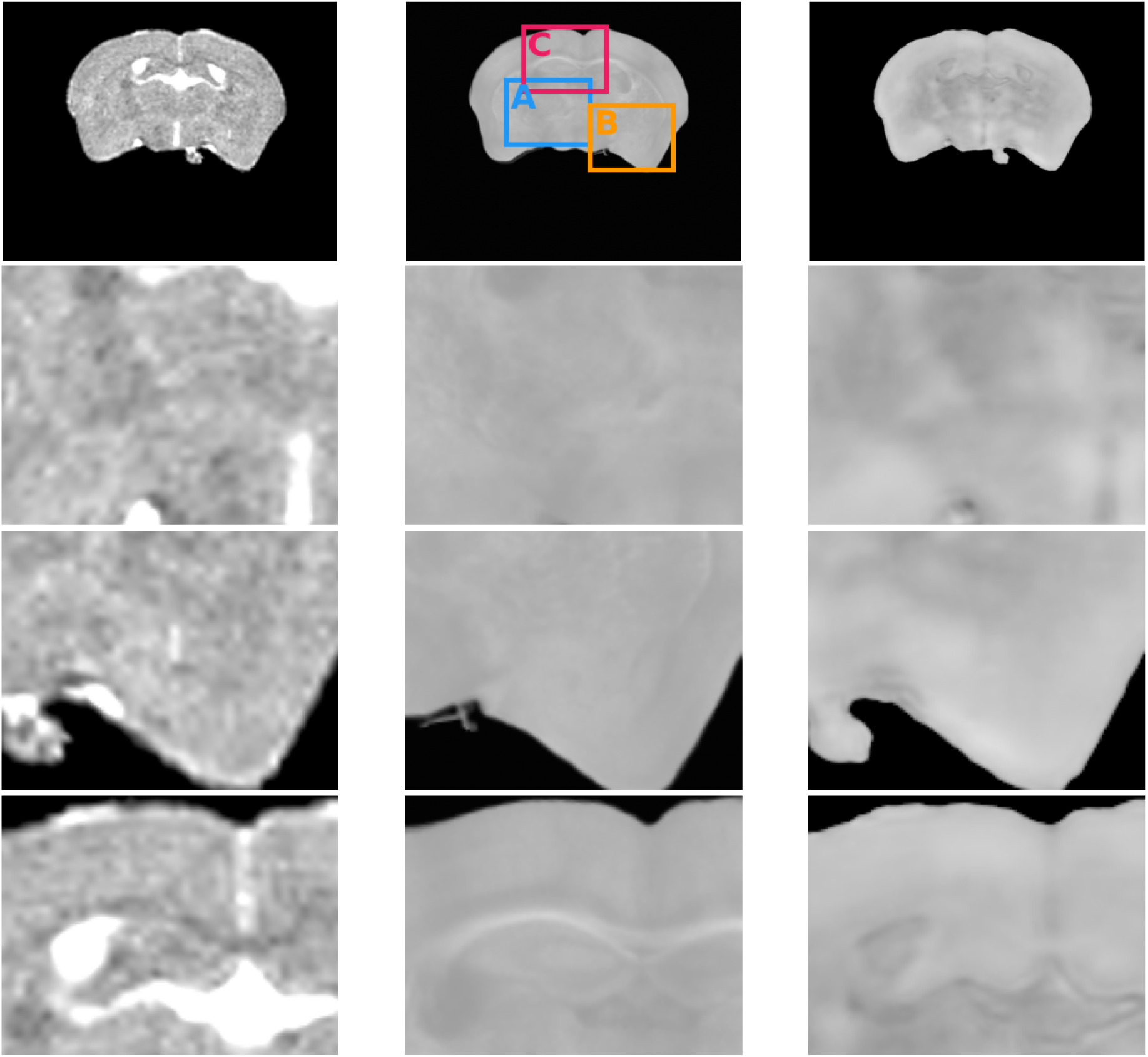
Coronal plane of one held-out subject. Left to right: the T2-weighted source, the PS-OCT reference with three regions marked (A, B, C) and magnified in the rows below, and the synthesised volume on the 4x grid. The source is shown at its native sampling and the reference at the reconstructed 10 µm spacing; the enlargement rows show the same physical field for each column. The outline and the hippocampal region are reproduced without visible grid structure. The reference contains sectioning marks and dark lines that the synthesis smooths rather than reproduces, which is the expected behaviour of a model operating inside the boundary rather than a defect to be corrected.

Across the nine leave-one-subject-out folds, the model was evaluated on parenchymal voxels of the union brain mask. It reached a peak signal-to-noise ratio of 22.01 *±* 0.95 *dB* (median 21.75), a structural similarity of 0.7630 *±* 0.0258, and a learned perceptual distance of 0.1873 *±* 0.0226. We computed the perceptual metric on axial, coronal and sagittal slices at five-voxel spacing, with scores aggregated by tissue-voxel weight. The methods state the three formulations and the reporting conventions behind these numbers; the identity of the feature backbone matters to the third value, and it is named there.

We report these numbers as an existence check, not as a comparison, and the low dispersion across folds is consistency, not accuracy. The single inference plane shown in Figure 3 carries the same standing: it is one plane of one held-out subject, not a survey of the cohort.

## 4 Discussion

This study asked whether a 100 µm MRI volume still carries information about a finer band when it is asked to supply that band. It answered with two readouts on a shared band index, evaluated at six candidate magnifications on nine paired mouse brains before any generator was trained. The finding worth carrying away is not the number four but the separation behind it. Geometric guidance and band energy fail at different candidates, so the boundary rests on two conditions and not on a single declining score. At 4x, a target voxel of 25 µm, both still hold, with the orientation readout at 0.661 against an analytic level of 0.50 and the energy readout at 0.438. That changes the status of the magnification itself. A magnification has been treated as a setting a model’s capacity is made to fill. It can instead be measured as a property of how much of the finer band the source still determines, at a stated baseline and independently of which learner is used. A study that improves one of the two readouts without moving the other has not moved that measurement.

### 4.1 Two readouts that fail at different candidates

The separation is the finding the argument turns on, and its mechanism is the one already established for the two estimators. Orientation comes from the principal eigenvector of a structure tensor, is insensitive to overall intensity, and survives where boundary geometry still constrains the finer band. That estimator is the one standard for serial optical coherence data [29], and its accuracy in brain-tissue microscopy depends on window scale and noise in a way that has been measured directly [30]. The comparison of optical and diffusion-based orientation has been reported at a single fixed scale, usually for a reconstructed fibre angle, in the human medulla [4], in post-mortem validation frameworks for diffusion MRI [5], and in the human brainstem [6], while the bridging of white-matter geometry across anatomical length scales has been treated separately [7]. None of those works asks how the agreement behaves as the target band is refined, and this work asks that question across a ladder of target bands. Energy comes from a smoothed scalar field whose passband cuts into the finest bands, so it loses explanatory power earlier, and part of that loss is a property of the estimator rather than of the tissue. Across the six candidates the geometric excess over the isotropic level and the explained variance move in opposite directions, and the two cross instead of declining together. The boundary inherits both behaviours. The geometry makes 4x interpretable, since a source-supplied direction is still there to stand on, and the energy makes it a boundary, since past that candidate the predictability of the band falls away faster than the direction does.

The condition under which that reading holds is narrow and worth stating in full. It requires a target that supplies both an orientation-bearing quantity and a phase-invariant energy from the same acquisition, which is the property that makes a polarisation-sensitive measurement usable as a structure reference [2, 3], and that a contrast carrying only one of the two cannot supply. It requires a source whose grid is isotropic so that a band defined on frequency modulus means the same thing on every axis, together with a tensor and an envelope computed identically on both sides. That requirement is also why the accuracy of the tensor has to be reported through its window and noise settings rather than assumed [30]. It requires, finally, that the bands be read as answers to different questions and not as one axis of difficulty, since each candidate interrogates a different physical period and a per-band change therefore mixes a loss of source information with a change in what is being asked. A pair that meets the earlier conditions but not this last one will produce the same two curves and cannot claim the same interpretation from them.

The practical consequence follows from the same asymmetry. A synthetic volume presented at 6.25x is not a numerical error. The super-resolution literature has treated the magnification as a parameter of the output representation, and not as a quantity the data certify: from adversarial and perceptual objectives [20, 21] through diffusion-based and implicit-representation formulations [22, 23] and the surveys that organise them [24]. Frequency has had a different job in that literature, serving as a target instead of a measuring rod. Spectral terms penalise the distortion that pixel losses under-weight [25], and frequency-selective restoration couples a frequency branch to a spatial one [26]. The distinction between more pixels and more resolving ability has been argued explicitly for deep-learning super-resolution in microscopy [27], in a domain where the answer decides whether an image can be trusted. At 6.25x the geometric excess over the isotropic level has fallen to 0.053, and the energy readout explains a third of the variance, so the source still partly carries position and orientation while the finer detail is not. Past that point the evidence responsibility moves. A band whose detail is produced and not recovered is not wrong, but the burden of showing that its detail is informative, and not merely plausible, shifts to whoever presents it.

### 4.2 Strengths built into the construction

Three methodological strengths follow from how the probe is built and not from its results. The measurement needs no trained generator, because the two readouts are computed from tensors and from band energies on a frozen mapping between source and target. Its answer therefore does not move when an architecture or a loss is changed, and it stands as a ceiling on what any particular learner can achieve. This is what separates it from the family of designs that made cross-domain mapping routine: paired image-to-image translation [10], cycle-consistent learning without pairs [11], CT and MR synthesis [13, 14], super-resolution within a single modality [15], and the surveys that order these designs by architectural family [16, 17]. Diffusion models have since supplied the generative step in medical synthesis [12], and reviews of generative medical imaging order the remaining designs [18, 19]. Each of those reports what a chosen model reaches at the scale it was trained on, so the number is a property of the model and the training set. The question asked here is what the source still determines at a scale it was not validated on, and that quantity is bounded above by the data, not by capacity. The probe also reports a baseline instead of an impression: the geometric readout is read against the analytic isotropic level, and the energy readout against a leave-subject-out score, so that agreement never rests on visual judgement. Its interpretive limits sit in the construction, because the shell integration keeps the band index away from the Nyquist edge, and the envelope width has a passband that is published with the score. A reader can therefore see which part of a decay belongs to the data and which to the instrument.

### 4.3 Three pipeline effects and their directions

The cost of building the readouts this way is that an analysis step, and not the modality pair, carries each one, and the three steps that matter have effects of known sign. The envelope kernel removes part of every band it scores, and the removed fraction grows with the candidate. It therefore pushes the energy readout down at the fine end and makes the decay steeper than the coupling alone would make it. The reference-side smoothing acts in the other direction. The reference was smoothed with a Gaussian of 12.5 µm, one half of the target voxel, before resampling to 25 µm. The 4x band therefore arrives attenuated across its upper half, with amplitude 0.74 at 10 cycles/mm, 0.50 at 15 cycles/mm and 0.29 at 20 cycles/mm. A partially smoothed target is easier to predict, so the *R*^2^ of 0.438 describes the target as represented and that value is pushed up, the more so because the half-amplitude point at 10.60 cycles/mm falls inside the 4x band instead of above it. The third has no fixed sign. The degradation cut-off is held at 5.0 cycles/mm for every candidate, which sits at the lower edge of the 2x band and one fifth of the way to that of the 10x band. Leakage into the finer bands is therefore suppressed less than leakage into the coarse ones, and any such leakage would raise *R*^2^ at the fine end and flatten the curve by an amount that depends on the spectrum of the MR volume. The three do not cancel, since they act on different quantities, and we therefore report the boundary as a property of this pipeline and not of the modality pair alone.

### 4.4 What transfers, and the rule that carries it

The criterion transfers because its inputs are generic: paired images of the two modalities, a registration that can be frozen, a null computable from geometry or from a permutation, and a linear predictor. What it returns is a frequency, not a class label, so a method operating inside the boundary is not certified as faithful; it is certified as operating in a band where the source still constrains something. The value four does not transfer. It should be expected to move with a different fixation protocol, a different reconstruction chain that changes the effective point spread, or a species whose tissue anisotropy has a different characteristic scale. A transfer should also not assume that the two readouts turn at the same candidate, since whether they do is precisely what the procedure measures.

What the margin enumeration adds is a division of labour a reader can carry to such a case. The geometric channel decides whether the fine candidates are excluded at all, and the energy channel decides whether the boundary moves down from 4x. A transfer that finds the geometry weak in every band should expect an empty answer and not a small boundary. One that finds the energy strong everywhere should expect the boundary to be set by geometry alone. Neither case is a failure, and each is a reading of the probe.

### 4.5 Limitations and next steps

Three limitations follow from the design of the probe and not from its execution. First, the readouts are pooled point estimates, and the pooling is what makes them comparable across candidates, since a per-subject summary would rest on a different voxel population in each band. What the pooling leaves unstated is whether the geometric excess at 4x is carried by most brains or by a subset, a question a per-subject decomposition would answer alongside the pooled medians without changing the criterion. Second, the six candidates interrogate different structures, from 200 to 500 µm at 1x to 20 to 40 µm at 10x, so the fall in explained variance from 0.627 to 0.175 mixes a loss of source information with a change in what is asked, and dividing the two contributions requires a reference sampled more finely than the 10 µm of this cohort. Third, the tensor medians rest on one anisotropy cut at 0.3, whose admitted share falls from 98.5 to 78.3 per cent across the candidates, and sweeping that cut is a recomputation over stored volumes that needs no new data. These limitations bound how tightly the crossing can be placed and leave where it lies untouched: a reference sampled more finely than the one used here is the addition that would divide the decline between the source and the estimator, while the per-subject decomposition and the threshold sweep would fix the width of the interval around 4x.

## 5 Conclusion

Whether an MR volume can be enlarged to a given grid is a question about its content, and this paper answers it before any network is involved. On nine paired mouse brains a probe built on a shared band index tracked that content from a period of 200 to 500 µm down to 20 to 40 µm. The median absolute cosine between macro-scale and micro-scale principal directions stayed above the isotropic level of 0.50 until 4x and then fell to 0.518 at 10x, while the leave-subject-out *R*^2^ of the band envelope fell monotonically from 0.627 to 0.175. The two curves cross at 4x, a target voxel of 25 µm, and that candidate is the last at which both conditions still hold. Beyond it at least one fails, so detail at those bands is produced and not recovered. The worth of the result lies less in the number than in the instrument that yields it: a shell decomposition, a phase-invariant envelope and a tensor comparison against an analytic null. That construction outputs a frequency, its inputs are a paired reference and a registration that can be frozen, and its verdict does not change when the band axis is normalised either way. It lets a user of a super-resolution method state, for a source and a reference they actually hold, how far the enlargement stays anchored in the data, and it moves that statement from the side of the model to the side of the measurement.

## Acknowledgements

This work was supported by the Brain Science and Brain-like Intelligence Technology project of the National Science and Technology Major Project of China (2021ZD0200201).

